# Identification and prediction of G-quadruplex RNA-binding proteins with roles in transcription and phase separation

**DOI:** 10.1101/2023.04.01.535204

**Authors:** Johanna Luige, Alexandros Armaos, Gian Gaetano Tartaglia, Ulf Andersson Vang Ørom

**Author notes:** Co-corresponding authors, Correspondence to (GGT) or (UAVØ). These authors contributed equally (Johanna Luige, Alexandros Armaos).

## Abstract

RNA-binding proteins are central for many biological processes and their large-scale identification has demonstrated a broad range of functions. RNA G-quadruplexes are important regulatory elements occurring in both coding and non-coding transcripts, yet our knowledge of their structure-based interactions is limited. Here starting from theoretical prediictions, we show experimentally that a large number of chromatin-binding proteins bind to RNA G-quadruplexes and we classify these based on their RNA G-quadruplex-binding potential. Combining experimental identification of nuclear RNA G-quadruplex-binding proteins with computational analysis, we create a prediction tool that can assign probability score for a protein that it binds RNA G-quadruplexes. We show that predicted G-quadruplex RNA-binding proteins exhibit high degree of protein disorder and hydrophilicity, and suggest involvement in both transcription and phase-separation into membrane-less organelles, particularly the nucleolus. Finally, we present this tool as a web application for estimating RNA G4-binding propensity for proteins of interest at http://service.tartaglialab.com/new_submission/clever_G4_classifier.

## Introduction

RNA have regulatory and structural roles in all cellular processes that are executed through RNA–protein interactions (1). The vast interconnection between RNAs and protein factors is reflected in the coordinated cellular responses to external signals or insults. This includes the regulation of transcription, where the interplay of RNA and protein factors controls assembly of the transcriptional machinery at enhancers and promoters (2). Accordingly, a growing number of dual specificity DNA–RNA-binding proteins (DRBPs) have been identified (3), and is a topic of active investigation.

The past decade has provided tremendous insight into RNA-binding proteins by the invention of interactome studies based on poly(A) capture and identification of binding proteins by mass spectrometry (4,5). Interactome-wide identification has been extended to subcellular compartments (3) and to refined protocols to purify all RNAs independent of poly(A) tails (1). A recent summary of RNA-binding proteins identified by various variations of interactome capture shows that 6,570 proteins have been proposed to bind RNA in some context either in cell culture or tissues (6).

G-quadruplexes (G4) are higher-order nucleic acid structures that form in guanine-rich sequences. The basis for G4 folding is the ability to form hydrogen bonds between two non-adjacent guanines, creating the G-quartet. As a result, the single-stranded nucleic acids can fold into four-stranded G-quadruplexes, formed by stacking of two or more G-quartets. G-quadruplexes are stabilized by intercalation of monovalent cations between the G-quartets, enhancing the base-stacking interaction.

RNA G4s are dynamic structures and their function is believed to be regulated by RNA-binding proteins (7), and interaction with RNA G4s can mediate both competition and cooperation between RNA-binding proteins (8). *In vitro*, the folding of G4 structures is affected by cations in the buffer, where for RNA potassium (K^+^) stabilizes G4 formation the most, sodium (Na^+^) provides intermediate stabilization and lithium (Li^+^) provides the least stability, which can be exploited in experimental setups to determine G4 function and identify G4-RNA-binding proteins (G4RBP) (9). Additionally, G4 formation at the boundaries of topologically associated domains indicates their crucial role in three-dimensional chromatin organization, that is known also to involve the architectural proteins CTCF (10) and YY1 (11).

There have been multiple screening experiments for identifying G4RBPs, and most of these data are incorporated into the recent database QUADRatlas (7) giving information on transcripts with G4-forming RNA sequences and G4RBPs. As an example, nucleolin is an exemplary G4RBP, that has been shown to bind to the mRNA and promoter of VEGFA to regulate its transcription (12).

In our previous work, we find that the nuclear RNA-protein interactome reveals new RNA-DNA dual binding proteins with an involvement in the DNA damage response (3), linking DNA and RNA binding properties to a number of proteins. In addition, a recent study proposes general importance for RNA in mediating transcription factor function (2). Here, we expand the RNA-protein interactome by exploring the combination of experimental and computational approaches to identify G4RBPs, demonstrating physicochemical properties involved in specifying protein binding to G4 RNA. We apply this on our webserver to predict proteins binding to G4 RNA, and propose a function in phase-separation of G4RBPs, especially in the nucleolus.

## Results

### Chromatin-associated RBPs often bind to G4 RNA structures

Given the impact of RNA on transcription factor function (2) and the number of dual DRBPs identified in interactome capture studies (3,6), we wondered whether RBPs associated with chromatin are generally able to directly bind to G4. To predict the RBP interaction preference for G4, we exploited the *cat*RAPID approach [**Materials and Methods;** (13)] that computes the interaction propensity using secondary structure properties, van der Waals and hydrogen bonding potentials. In our calculations we built two models of secondary structure occupancy corresponding to folded (structured) and unfolded (linear) G4 RNA. We used an optimized G4 forming sequence, G4A4, consisting of repeats of 4 Guanosines followed by 4 Adenosines (see **Materials and Methods**) and computed the interaction propensity for 283 chromatin-related proteins identified in K-562 nuclei from previous work (3). We calculated G4A4 contacts for each protein (14) and found that 182 of them bind preferably to folded G4A4 (with K^+^), while the rest interacts with unfolded G4A4 (with Li^+^) (**Figure 1, Supplementary Table 1; Materials and Methods**), suggesting a widespread binding preference of chromatin-associated RBPs to G4 RNA.

**Fig. 1.**
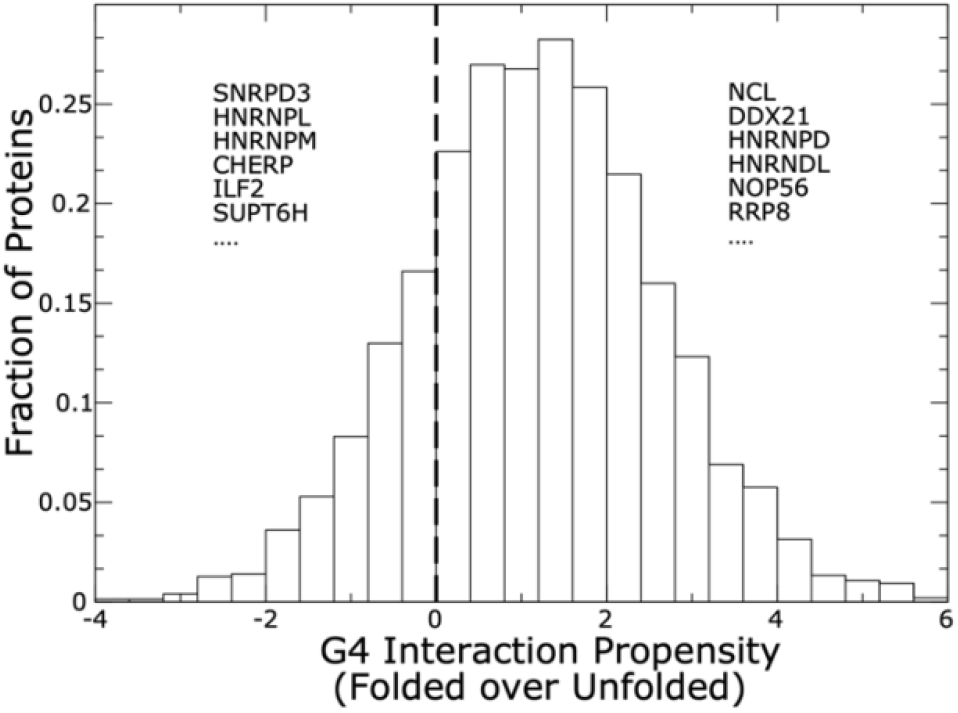
Chromatin RBPs are predicted to bind to the G quadruplex structure. *cat*RAPID predictions of RBP interactions with folded (structured) and unfolded (linear) G4A4 indicate that the majority of chromatin-related proteins have a binding preference for folded G4A4. Examples of proteins with preference for folded and unfolded G4A4 are reported.

### Experimental identification of G4RBPs

Next, we proceeded to experimentally assess the binding preference of RBP through an *in vitro* approach. In our experiments, we used the G4A4 oligo as used for computational modeling and coupled it to biotin to allow for purification of bound proteins from K-562 nuclear extract. We performed the experiment in triplicate in the presence of K^+^ or Li^+^ cations to affect the stability of G quadruplex structures in the G4A4 oligoribonucleotide. We show with circular dichroism that the RNA G4A4 oligo folds into a parallel G4 (**Figure 2a**), as would be expected for a canonical G4-forming RNA (15). To set up the conditions and demonstrate sensitivity of our approach to purify G4RBPs we used the well-characterized G4 RNA binder NCL. The protocol for protein purification is shown as a diagram in Supplementary Figure 1. For NCL we show binding to G4A4 by western blot in **Figure 2b** and that this binding is significantly stronger in the presence of K^+^ compared to when Li^+^ is used in the assay buffer, in agreement with the known impact of cations on G4 RNA structure stability (9). We then subjected samples to LC-MS/MS mass spectrometry in triplicate and identified proteins that were significantly enriched or depleted in K^+^ buffer compared to Li^+^ buffer (**Figure 2c, d)**. By mass spectrometry we detect a total of 1,204 proteins of which 151 and 83 are enriched and depleted, respectively, in the purification using K^+^ cations in the binding buffer (Supplementary Table 2).

**Figure 2.**
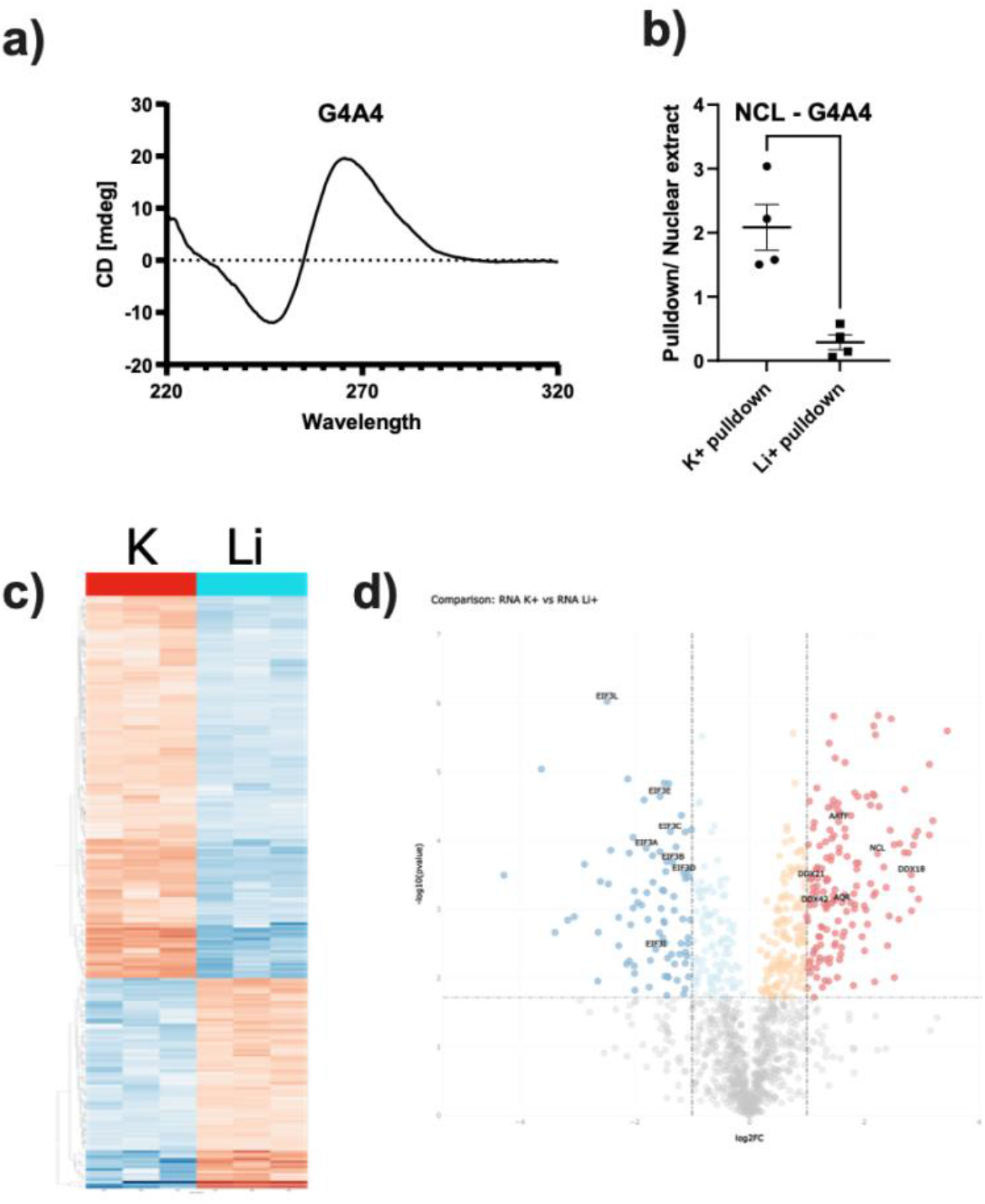
Experimental identification of G4-RBPs. A) Circular dichroism specter for G4A4 RNA. B) Pulldown of NCL from nuclear extract is dependent on folded G4 (K^+^), n=4, statistical significance level determined by Student’s t-test (*p<0,05). C) Hierachical clustering of K^+^ and Li^+^ enriched proteins across three replicate pulldown experiments. D) Volcano plot for significantly enriched proteins. G4BPs AATF, DDX21, DDX42, AQR and NCL are shown.

### G4RBPs are generally known RBPs

Of the 151 proteins with increased binding upon G4 stabilization 83 (55 per cent) have a previous annotation as G4RBP according to QUADRatlas (7), while for the proteins with decreased binding upon G4 stablization 26 (31 per cent) are classified as G4RBP by QUADRatlas (**Figure 3a, Supplementary Table 3**), demonstrating that our assay enriches for *bona fide* G4RBPs. Of the 68 proteins not previously shown to associate with G4 RNA, 19 have the GO-term DNA binding (p=0.00029), including CTCF and TOP1, pointing towards G4-binding as a bridge between DNA and RNA binding properties for chromatin-associated proteins. At the same time, this comparison shows that several aspects of proteins and G4-forming RNA sequences are likely to impact their binding. When assessing Gene Ontology (GO) for the entire set of identified proteins we see that the overall identified proteins in the mass spectrometry analysis regardless of response to K^+^ and Li^+^ show 48.3 percent with the GO term RNA binding (566 of 1,172 with annotated GO term) (**Figure 3a**), supporting the quality of our *in vitro* pull-down assay. For proteins that are enriched in K^+^ buffer this percentage increases to 79.9 (119 of 149 with annotated GO term) and for proteins decreased in K^+^ buffer compared to Li^+^ buffer this percentage is decreased to 22.9 (**Figure 3b**). These data show that the proteins with increased binding in K^+^ buffer are *bona fide* RNA binding proteins where most have been annotated with the GO term RNA binding, supporting G4 RNA binding as a central property for RBPs. In contrast, proteins that preferentially bind the G4A4 RNA in Li^+^ buffer are non-canonical RBPs. Overlap with the nuclear interactome from K-562 cells (3) shows that 48 of the proteins increasing binding in the K^+^ buffer overlaps with the previously annotated K-562 nuclear interactome, whereas only a single protein increasing binding in Li^+^ buffer does, showing that responsiveness of proteins to K^+^ for binding to stabilized G4 RNA sequences supports an annotation as true RBPs. In total, 229 of the identified proteins overlap with the 343 proteins (67.8 per cent) in the K-562 nuclear interactome. 766 proteins neither overlap with the K-562 nuclear interactome nor change their binding in response to K^+^ and Li^+^, and amongst these 254 (out of 761 (33 per cent) with annotated GO term) have the GO term RNA binding, suggesting that these might predominantly consist of background due to the *in vitro* nature of the assay. Of the 31 proteins (30 with annotated GO term) not annotated as RNA-binding that are enriched in K^+^ buffer, 12 have the GO term transcription, and 12 have the GO term DNA binding with a substantial overlap between the two groups (**Figure 3c**). In the K-562 nuclear interactome not overlapping with the proteins identified in this study there is no significant enrichment for DNA-binding proteins or transcription factors, suggesting that G4RBPs does have an important role for connecting RNA- and DNA-binding proteins (**Figure 3d**).

**Figure 3.**
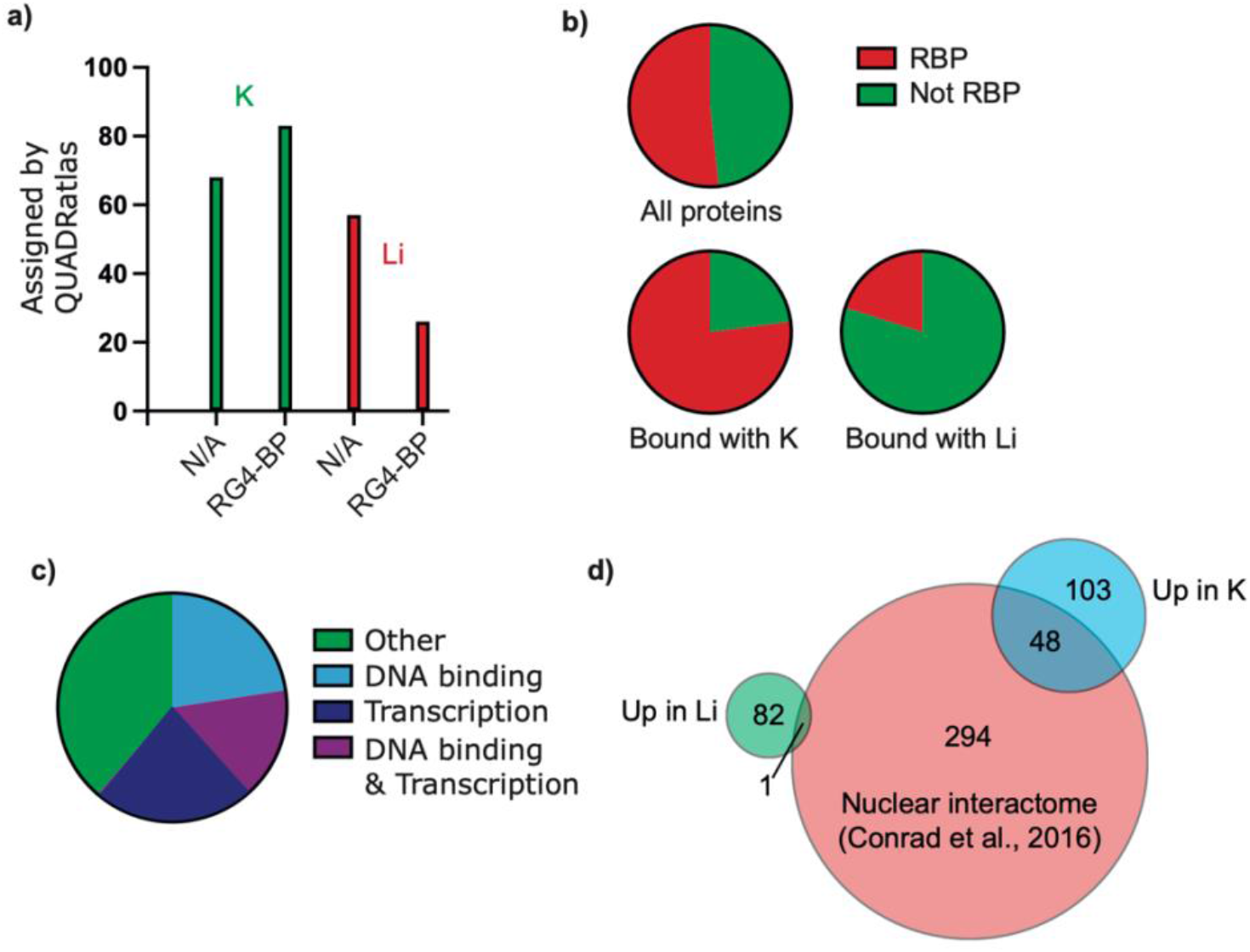
Characterization of proteins identified in G4A4 pull-down. a) Comparison to QUADRatlas of the proteins identified with pull-down and mass spectrometry. Graph shows how many proteins for each K and Li pull-down conditions that have been assigned as G4RBP (RG4-BP) or has not been shown to bind to G4 RNA sequences (N/A). b) Analysis of GO term RNA-binding protein (RBP) for identified proteins in the three groups all proteins (1,172), Bound with K (149) and Bound with Li (81) c) GO-analysis of proteins bound in K not annotated as RBP. d) Overlap with NIC. A Venn diagram showing the overlap between RBPs identified by nuclear interactome capture and by binding to the G4A4 sequence in a stabilized (Up in K^+^) or destabilized (Down in K^+^) state.

### Computational characterization of G4RBPs

Having established that proteins enriched in K^+^ buffer are *bona fide* RBPs enriched in conditions where G4A4 is stabilized in a G4 structure, we set out to explore the properties of these protein groups (*i*.*e*. enriched in presence of K^+^ or Li^+^).

We assessed the performance of our computational tool *cat*RAPID (16) on the experimentally identified G4RBPs (**Supplementary Table 4**). *cat*RAPID predictions agree with our experimental results, with interactions for folded G4A4 preferably found in presence of K^+^ ions and interaction for unfolded G4A4 observed in presence of Li^+^ ions (**Figure 4a**). Specifically, we calculated how many times the folded state is preferred over the unfolded one for the different RBPs at different *cat*RAPID interaction propensities. The analysis was performed for the K^+^ and Li^+^ ions group independently.

**Figure 4.**
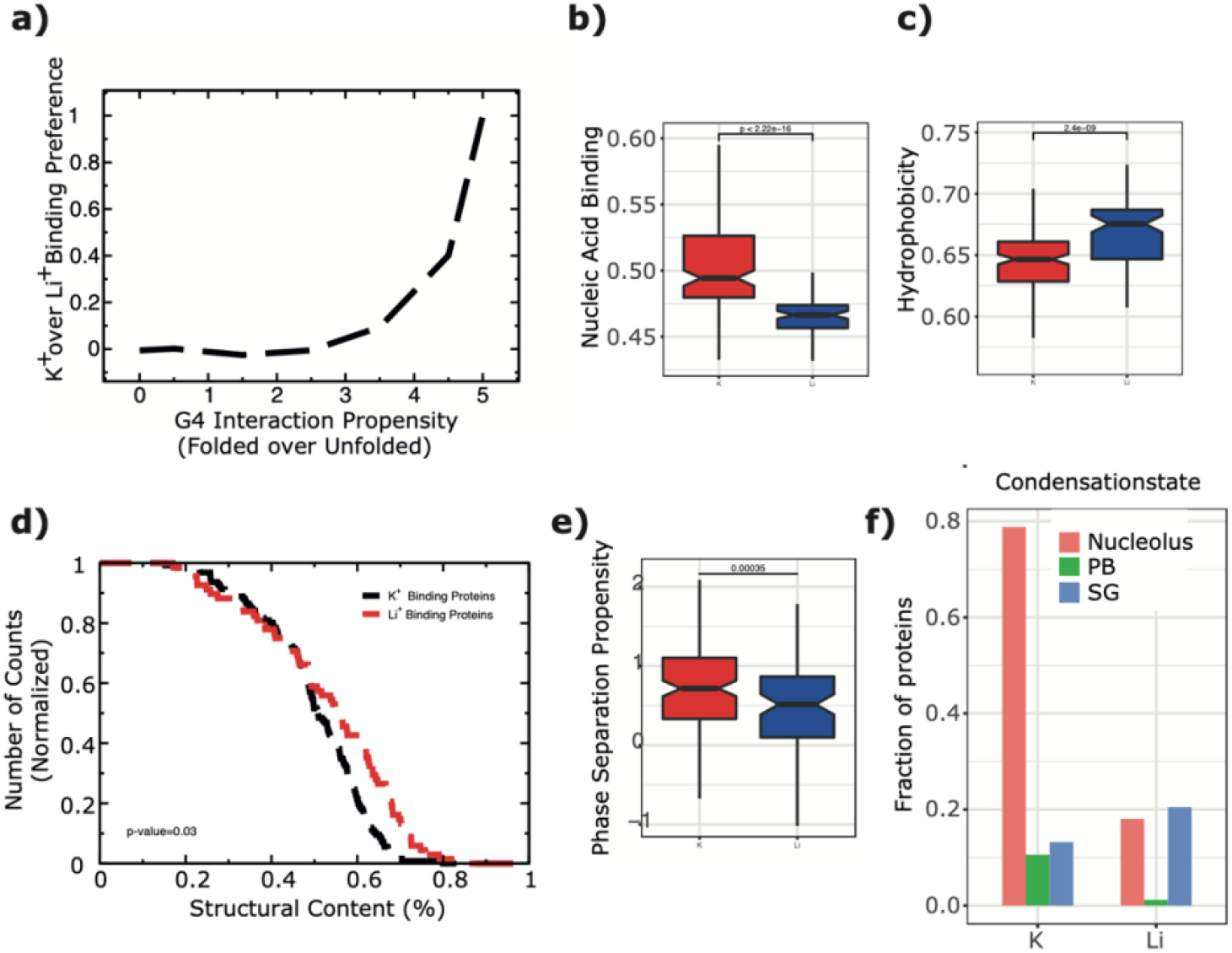
**a**) Experimentally-validated interactions in presence of K^+^ are predicted by *cat*RAPID to preferably bind folded G4. **b)** Proteins interacting with G4A4 in presence of K^+^ ions are predicted to be canonical RBPs (18), and **c**) depleted in burial (19). **d)** Structural content in K+ and Li+ protein groups as determined by Alphafold. e) Phase separation propensity as calculated with *cat*GRANULE. f) Condensation state of proteins interacting with G4A4 preferentially with K+ or Li+.

For further insight into the properties of G4RBPs we used the *clever*MACHINE algorithm that discriminates between two protein datasets using physico-chemical properties contained in their primary structures. The first dataset contains a signal (protein interacting with G4A4 in presence of Li^+^ ions) and the second is compared to it (proteins interacting with G4A4 in presence of K^+^ ions) (17). The algorithm uses physico-chemical propensities derived from experimental data **(Materials and Methods**) and allows us to discriminate between the two groups with 96% confidence, with proteins identified in presence of K^+^ ions enriched in non-canonical RBPs (**Figure 4b**) (18) and depleted in burial, suggesting larger amount of structural disorder (**Figure 4c**) (19). To back up our predictions, we used AlphaFold to investigate the amount of structural content in the proteins sets and indeed found that K^+^ binding proteins are significantly enriched in disorder (**Figure 4d, Supplementary Table 5**).

### Disordered G4RBPs with a predicted role in phase separation

Disordered protein domains are known for their multifaceted roles in the formation of biomolecular condensates, in which protein and nucleic acids within a solution can dynamically undergo demixing, resulting in separation into distinct phases with different molecular composition (20). We investigated phase-separation ability of proteins using *cat*GRANULE, and found that proteins binding to G4A4 in the presence of K+ have indeed higher phase separation propensity (**Figure 4e; Materials and Methods, Supplementary Table 6**). We complemented our predictions with an analysis of protein occurrence in phase separated organelles such as nucleoli, stress granules and p-bodies (**Figure 4f, Supplementary Table 7**). In agreement with our predictions, proteins binding to G4A4 in the presence of K^+^ show enrichment across condensation states, especially nucleolar proteins, thus confirming *cat*GRANULE calculations. Recent studies suggest G-quadruplexes can drive phase separation (21,22), particularly in nucleolus (25,26), by allowing multivalent interactions through potential stacking of multiple folded G4s (23), as well as providing spacial recognition surfaces for protein partners (24).

### Computational identification of novel G4RBPs

To expand our data and to address the cellular repertoire of DRBPs we assigned G4-interaction propensity to all proteins with the GO term chromatin binding based on our findings for experimentally identified G4RBPs. We again used the *clever*MACHINE to calculate interaction propensities for chromatin proteins and classified them as Li^+^ and K^+^, with a lower cut-off for scoring as G4RBP set at 50 (**Supplementary Table 8; Materials and Methods**), assigning G4RBP properties (**Figure 5a, b**). In addition, *cat*GRANULE indicates enrichment of predicted K^+^ binding proteins, in agreement with our results from experimental datasets (**Figure 4f**).

**Figure 5.**
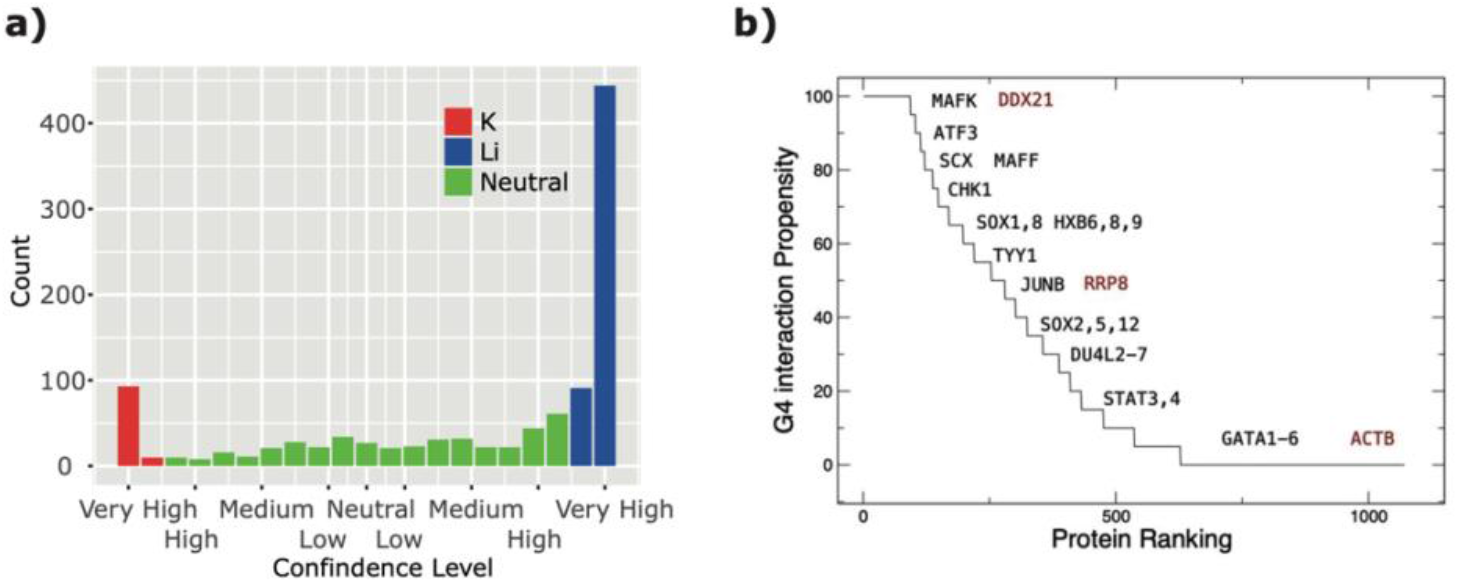
G4 interaction propensities of chromatin proteins. a) The number of chromatin proteins assigned as N and P (binding and non-binding to G4A4) by Clevermachine for each confidence level. b) The interaction propensity. Black is used to highlight examples of predicted classes. Red indicates controls present in our experimental data. The *clever*Machine was used for the calculations.

Having found that specific pysicochemical properties are signatures of proteins binding to G4 RNA we addressed the presence of protein motifs. Using PFAM database (Enrichr) we find that DEAD/DEAH box proteins, Helicase C and bromodomains are the most significantly enriched motifs in the proteins binding to G4 RNA in the presence of K (**Table 1**), whereas for the proteins with decreased binding in K buffer PCI (Proteasome component) and Tubulin domains are the most significantly enriched (**Table 1**).

**Table 1.**
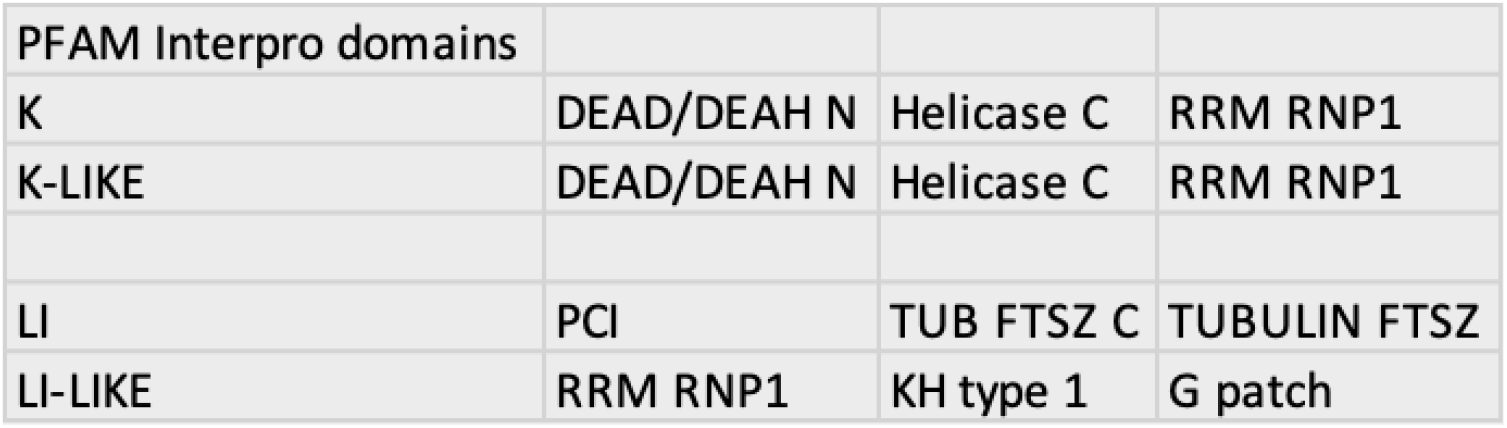
Pfam Interpro domains in identified and predicted G4-RBPs.

Comparing *clever*MACHINE-predicted K-like proteins and enriched domains, we find the same groups as for the experimentally identified, supporting DEAD/DEAH and Helicase C domains as important for binding RNA G-quadruplexes. The predicted Li-like proteins do not recapitulate experimentally identified proteins on the domain level, suggesting that no specific enrichment of protein domains repelling G4 RNA exist in our dataset. Of interest, the Li-like proteins are enriched for the RRM RNP1 domain as we also see in both K groups, suggesting that this domain is non-specifically enriched due to other factors than G4 RNA binding.

### Experimental validation of new predicted G4RBPs

To validate whether the candidates we assign as G4RBP bind to G4 RNA sequences within the cell, we used native RNA immunoprecipitation of endogenous candidate proteins that we predict as G4RBP, Chk1 and Ruvbl2, followed by RT-qPCR analysis of protein-bound RNAs (Figure 6a, b). We use *VEGFA, MYC, BCL-2* and *NRAS* as reference for endogenous G4 RNA, as these are some of the most thoroughly validated and prominent cellular mRNAs harboring 5’UTR G-quadruplexes (27–30). The negative control RNAs RN7SK and GAS5 are chosen for their apparent lack of annotated and predicted G4s according to the QUADRatlas, and being highly abundant transcripts.

**Figure 6.**
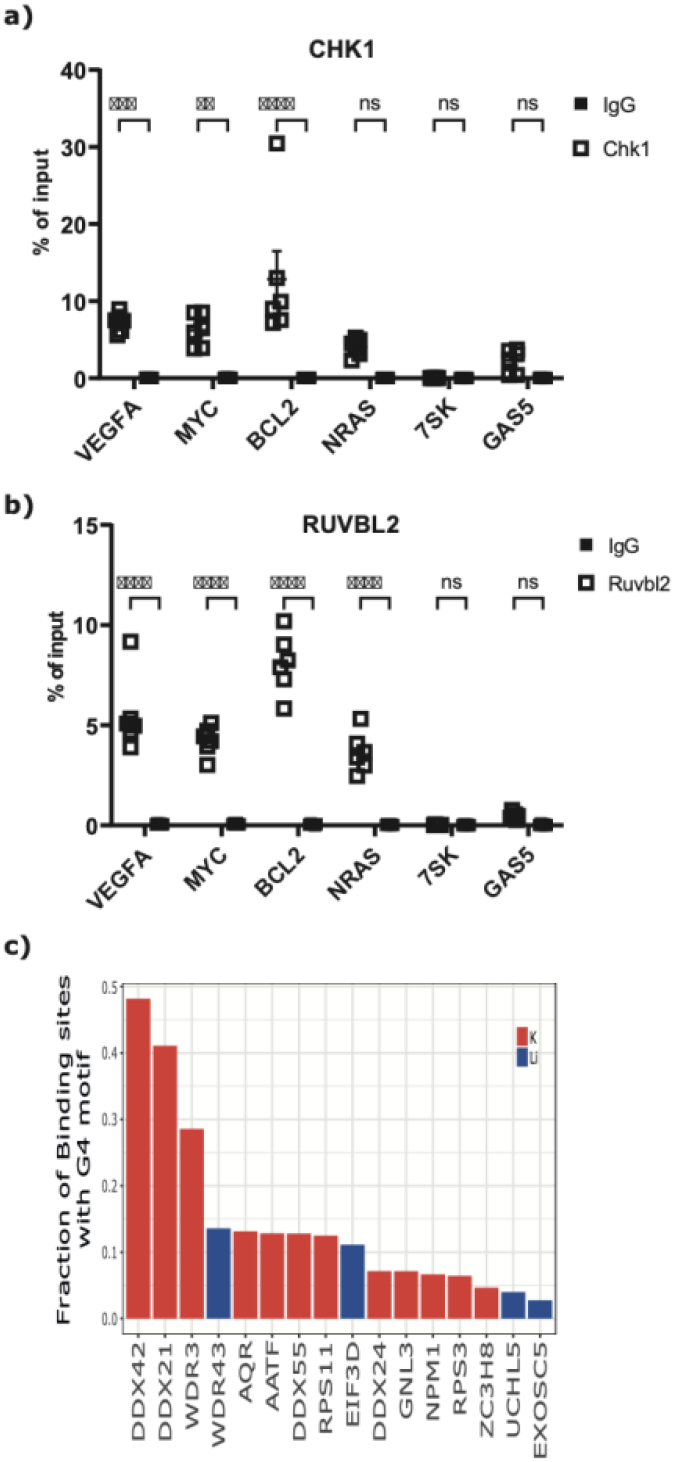
Validation of G4RBP interaction with RG4. a) Native RIP-qPCR experiment for Chk1 and Ruvbl2, validating RG4BP interaction with *VEGFA, MYC, BCL-2, NRAS* mRNAs, *7SK* and *GAS5* RNAs as control. b) eCLIP data for binding of candidate proteins at G4-forming sites in mRNA.

As a more general validation of the G4RBPs identified and predicted we used available eCLIP data to address the binding of 16 proteins to G4-sequences within their binding sites at RNA (**Figure 6c**), showing a preference of K-enriched proteins to bind within G4 forming sequences at RNA.

## Discussion

RBPs have been investigated at large scale in the last decade following the invention of poly(A) mRNA interactome capture (4,5). Approaches to identify proteins binding to all RNA has subsequently been developed (1). Common to these existing approaches is that they do not consider RNA structural elements such as G-quadruplexes, that are context-dependent and responsive to *e*.*g*. cations (9) and stress (31). In this study, we show by experimental large-scale identification of G4RBPs that several proteins previously not detected to bind RNA in K-562 nuclei (3) are RNA binding, and use computational tools to expand to the hypothetical complete G4RBP proteome. DRBPs have been identified by serial interactome capture of the cell nucleus to be a poly(A) mRNA enriched class of proteins (3) suggesting an intimate relationship between DNA- and RNA-binding properties for transcription factors and DNA damage proteins, where RNA could facilitate DNA binding and modulate enzymatic activity, as recently suggested to be a general feature for transcription factors (2). G4 forming sequences are present particularly at enhancers, promoters and within 5’UTR encoding sequences, and are thus well-positioned in the genome and transcriptome to link DNA- and RNA-binding properties for specific groups of proteins.

In addition, we show that G4RBPs accumulate in membrane-less organelles as the nucleolus and p-bodies, where binding of G4RBPs to mRNAs could mediate an efficient regulation of translation and localization of regulatory intracellular bodies. Indeed, through high-throughput dimethylsulfate probing it has been shown that the G4 structure forms upon stress (32) and several reports indicate that G4 sequences can induce phase separation (8,21). We note that the proteomes of nucleolus and p-bodies are characterized by proteins with higher degree of intrinsically disordered regions (IDRs), as well as well-known RNA-binding domains, such as RRG/RG motif (34), RRM and DEAD domains (34,35). Studies of DDX ATP-dependent helicases in multiple species show their ability to induce phase separation through low complexity protein domains, determined by the ATP hydrolysis state and the resulting interactions with their RNA substrates (36). Most importantly, this property of DDX helicases allows for turnover of the membrane-less compartments and can facilitate RNA partitioning between different granules, both in the cytoplasm and nuclear environment.

Several interesting examples of G4RBPs result from our analysis, including the enhancer-binding protein CTCF that has been shown to bind G4 DNA (37) and to require RNA for recruitment to chromatin (10). We also find YY1 that is central to enhancer function and has been reported to bind G4 DNA (11). From these data it is tempting to speculate that G4 at transcribed enhancers could facilitate transcription factor recruitment with impact on both enhancer function and chromatin landscape. At another cellular level, we predict TOP1 to be a G4RBP. TOP1 has recently been shown to bind G4 at DNA (38) and facilitate transcription of the MYC gene, that encodes a well-characterized G4-containing mRNA.

Finally, we suggest 19 proteins involved in DNA damage to be G4RBPs, including CHK1. CHK1 is a cell-cycle checkpoint kinase that has also been shown to be involved in DNA double strand break repair (39). The recognition of transcribed G4 RNA at DNA damage sites could be one of the underlying aspects of RNA requirement for DNA damage repair that has been reported (40), in agreement with the formation of R-loops at DNA break sites that are bound by the well-charaterized DEAD-box helicases (40), that we also see significantly enriched in our data.

In conclusion, we identify a set of bona-fide RBPs that recognize G4 RNA and use the biophysical properties of these proteins to model which chromatin-binding proteins have G4 RNA-binding propensity. We provide an overview of DRBPs that could connect DNA to RNA in biological processes as enhancer function, transcription and DNA damage repair; show evidence of G4 RNA-mediated localization of proteins to nucleolus and p-bodies; and present the biophysical properties and protein motifs important for recognition of RNA G4 structured sequences, that can be accessed as a tool on the webserver http://service.tartaglialab.com/new_submission/clever_G4_classifier.

## Conflict of interest

The authors declare no competing interests.

## Author contributions

JL and UAVØ conceived the experimental strategy and JL performed all experiments. AA and GGT conceived computational analysis strategies and performed computational analysis. All authors contributed to analysis and interpretation of the data. GGT and UAVØ drafted the manuscript and all authors commented on and accepted the final manuscript. GGT and UAVØ secured funding and supervised computational and experimental work.

## Ackowledgments

The mass spectrometry analysis was made by Proteomics Research Infrastructure at University of Copenhagen. Work in the author’s laboratories is supported by the ERC ASTRA_855923 and H2020 projects IASIS_727658 and INFORE_825080 (GGT), The Novo Nordisk Foundation, Independent Research Fund Denmark, The Lundbeck Foundation, Danish Cancer Society and Carlsberg Foundation (UAVØ).

---

**Supplementary Table 1**.

*cat*RAPID predictions of proteins binding to folded or linear G4A4. The **first column** contains gene name, **second column** corresponds to the Uniprot code, the **third and fourth columns** report the amino acids considered, **fifth and sixth columns** are the scores for the folded and linear G4A4, the **seventh column** is the average of the difference between columns sixth and seventh over all fragments of a protein, **eight column** is the maximal difference between columns sixth and seventh over all fragments of a protein.

**Supplementary Table 2**.

Detected proteins in all conditions in mass spectrometry.

**Supplementary Table 3**.

Proteins detected and differentially bound in mass spectrometry analysis, including QUADRatlas status.

**Supplementary Table 4**.

Chromatin RBPs from Conrad et al 2016 with prediction scores for binding to G4A4 in K^+^ and Li^+^ buffer.

**Supplementary Table 5**.

Secondary structure properties binding in presence of K^+^ and Li^+^ ions. **First column** is the AlphaFold name of the pdb. **Second, third, fourth, fifth columns** report the amount of Coil, Turn, Helix and Strand per protein. **Sixth column** is the protein length and **seventh column** the amount of structure normalized by length used in **Figure 4d**.

**Supplementary Table 6**.

Phase separation propensity.

**Supplementary Table 7**.

Condensation state.

**Supplementary Table 8**.

Proteome-wide chromatin-binding proteins with prediction scores for binding to G4A4 in K^+^ and Li^+^ buffer.

## Additional Data

The mass spectrometry proteomics data have been deposited to the ProteomeXchange Consortium via the PRIDE partner repository with the dataset identifier PXD041154.

## Material and Methods

### Tissue culture

K562 human myelogenous leukemia cell line (ATCC CCL-243) was obtained from ATCC. Cells were cultured in RPMI-1640 Glutamax (61870-036, Thermo Fisher Scientific) medium supplemented with 10% FBS (Gibco), 100 U/ml penicillin and 100 μg/ml streptomycin. K562 cells were maintained in 2 × 105 to 1 × 106 viable cells/ml.

### Circular dichroism spectroscopy

Circular dichroism for G4A4 RNA was carried out with Jasco J-810 spectropolarimeter. RNA oligonucleotide G4A4 (5’-AAAAAAGGGGAAAAGGGGAAAAGGGGAAAAGGGGAAAAAA-3’) was purchased from Merck. CD analysis of 2,5 µM RNA was carried out in the buffer used for G4-pulldown, containing 25 mM HEPES pH 7.0, 150 mM KCl, 5% glycerol. Spectral signatures were recorded over 220 to 320 nm wavelength.

### G-quadruplex pull-down

K562 cells were harvested and washed in PBS before lysis. Initial cytoplasmic lysis of K562 cells was carried out using an isotonic lysis buffer 0.1% Igepal CA-630, 150 mM NaCl, 50 mM TRIS pH 8.0, supplemented with protease inhibitor cocktail (P8340, SigmaAldrich) and 1 mM PMSF, incubating on ice for 10 minutes. Nuclei were collected by centrifugation for 3 min at 300 g. After removing the cytoplasmic fraction, nuclei were suspended in K+ or Li+-containing Buffer D (10 mM HEPES, 100 mM KCl/LiCl, 5% glycerol, 1mM PMSF, 1% protease inhibitor cocktail), incubated on ice for 10 min. 3 times 30 seconds ON/OFF sonication was carried out at high intensity (Covaris S2 Focus Ultrasonicator) to disrupt the chromatin and release nuclear proteins. Lysates were centrifuged for 30 minutes at 4 °C at 17 000 x g.

3’ biotinylation of G-quadruplex forming RNA oligonucleotides was carried out in 30 µl reaction, with 100 pmol RNA, 2 nmol biotinylated cytidine (19519016, Thermo Fisher Scientific), 2U T4 RNA ligase (EL0021, Thermo Fisher Scientific) in 1X ligase buffer, 20 U SUPERaseIN RNase inhibitor and 15% final concentration of PEG-8000. Ligation was incubated at 16 °C for 16 hours. Biotinylation reaction was cleaned with chloroform-isoamyl alcohol (24:1) extraction and ethanol precipitation. Pulldowns were carried out in either K+ or Li+ containing buffers (10 mM HEPES (pH 7.9), 150 mM KCl or LiCl, 5% glycerol, 0,2 U/µl SUPERaseIN), maintaining the buffer conditions throughout the experiment. Biotinylated RNAs were folded in K+/Li+ buffers by heating at 95 °C for 5 minutes, followed by incubation at room temperature for 1 hour.

Neutravidin-coated magnetic beads (Cytiva 78152104011150) were washed in K+/Li+ buffers 3 times 5 minutes with rotation before addition of the biotinylated G4-RNA. Each pulldown was carried out with 150 pmol biotinylated RNA, 50 µl beads, and 300 µg of K562 nuclear extract, incubating overnight on rotation at 4 °C. After incubation beads were washed with K+/Li+ pulldown buffer 3 times 10 minutes on rotation at 4 °C.

### Western blot

For western blot validation, elution of proteins was carried out by incubating the beads at 80 °C with 2.5X NuPAGE LDS sample buffer (NP0007, Thermo Fisher Scientific). Samples were loaded on 4-12% NuPAGE Bis-Tris polyacrylamide gel (NP0322, Thermo Fischer Scientific), running with NuPAGE MOPS SDS running buffer (NP0001, Thermo Fischer Scientific) at 110 V for 120 minutes. Transfer to Immobilon-P 0.45μm PVDF membrane (Merck Millipore) was carried out in NuPAGE Transfer buffer (NP00061, Thermo Fischer Scientific), supplemented with 10% methanol. Transfer conditions were 120 minutes with constant voltage at 100 V. Blots were blocked with 5% skim milk solution in PBS-T [1X PBS, 0.05% Tween-20]. Incubation with primary antibodies were carried out overnight as a 1:1000 dilution. Antibody incubations were followed by washing with PBS-T 3 times for 5-10 minutes. Secondary antibody against rabbit/mouse IgG was diluted 1:10000 in PBS-T. Signal was detected with the SuperSignal ECL reagent (34579, Thermo Fischer Scientific) and visualized with GE Amersham Imager 600. Western blot bands were quantified with Fiji software (Schindelin et al., 2012). Quantifired band intensities from pulldowns (n=4) were normalized to NCL levels in input lysates, and expressed as pulldown/nuclear extract ratio. Four replicate experiments were used for quantification, statistical significance was estimated with paired Student’s t-test (*p<0,05).

### Mass spec data analysis

All statistical analysis of LFQ derived protein expression data was performed using the automated analysis pipeline of the Clinical Knowledge Graph (Santos, A. et al. Clinical knowledge graph integrates proteomics data into clinical decision-making (41).

LFQ intensities were normalized by log2 transformation and proteins with less than at least 2 valid values in at least one group were filtered out.

Missing values were imputed using the MinProb approach (width=0.2 and shift=1.8, see (42).

Differentially expressed proteins were identified with unpaired t-test with Benjamini-Hochberg correction for multiple hypothesis (alpha=0.05, Fold Change (FC)=2) Hierarchical clustering was performed using all significant proteins. The selected proteins were then mean normalized.

### Native RNA-immunoprecipitation

Native RIP for endogenous mRNAs (*VEGFA, MYC, BCL-2, NRAS*) and non-coding RNAs (*RN7SK, GAS5*) was carried out using Chk1, Ruvbl2 and Rabbit IgG1 antibodies. Protein A and G coated Dynabeads (Thermo Fisher) were first washed in RIP lysis buffer, containing 25 mM TRIS, 150 mM KCl, 0.5% Igepal CA-630, 5 mM DTT, 20U/ml Rnase inhibitor (Rnasin, Promega), 1X Protease inhibitor cocktail (cOmplete, Roche). Prior to immunoprecipitation, antibodies were incubated with Protein A/G beads, using 4,8 µg of antibodies with 20 µl A+G beads. 20-30 million K562 were harvested for each RNA immunoprecipitation and lysed with 700 ul RIP lysis buffer for 25 min on ice, after which lysates were centrifuged 25 min at 4 degrees x 17 000g. 1% of lysate was removed for input analysis. RNA immunoprecipitation was carried out over 16 h at 4 degrees with rotation.

Beads were washed 5 times 10 minutes. Protein-bound RNA was eluted by incubating the beads with TRIzol, following RNA extraction. cDNA was prepared from equal volumes of immunoprecipitated and input RNA using Maxima H Minus reverse transcriptase (Thermo Fisher) and random hexamer primers. cDNA was diluted for detection of RN7SK transcript 1:130. qPCR analysis was carried out using Platinum SYBR Green (Thermo Fisher). CT values were converted by 2^^-CT method and normalized to input levels. Statistical significance was estimated using two-way ANOVA, with Šídák’s multiple comparisons test (GraphPad Prism). Statistical significance levels based on p-value: ns = not significant; *p <0.05; **p <0.01; ***p <0.001.

### G-quadruplex RNAs

[G4A4]4 AAAAAAGGGGAAAAGGGGAAAAGGGGAAAAGGGGAAAAAA

### Antibodies

CHEK1 (Abcam, EP691Y)

RUVBL2 (Abcam ab36569)

NCL (Abcam, ab136649)

Goat anti-mouse (Thermo Fischer Scientific, G-21040)

Goat anti-rabbit (Thermo Fischer Scientic, 31460)

### *cat*RAPID predictions of protein-RNA interactions

We employed the original version of *cat*RAPID (13) to predict G4 interaction propensity of chromatin, K+ and Li+ related proteins. The *cat*RAPID algorithm estimates the interaction through van der Waals, hydrogen bonding and secondary structure occupancy of both protein and RNA sequences (43). As reported in an analysis of about half a million of experimentally validated interactions (44), *cat*RAPID can separate interacting vs. non-interacting pairs with an area under the curve (AUC) receiver operating characteristic (ROC) curve of 0.78 (with false discovery rate (FDR) significantly below 0.25 when the Z-score values are >2). The secondary structure occupancy is defined by counting the number of contacts within the nucleotide chain. In our analysis, we used two types of secondary structure occupancy of RNA corresponding to the folded (structure) and unfolded (linear) G4A4. The 182 proteins preferably binding to folded G4A4 were identified considering that 75% of their contacts (polypeptide regions of 50 amino acids) have interaction propensities for folded G4A4 higher than interaction propensities for linear G4A4. The analysis is reported in **Supplementary Table 1**. Further information about the method can be found at http://s.tartaglialab.com/page/catrapid_group.

### *clever*MACHINE discrimination of protein sets

The *clever*MACHINE algorithm analyzes physico-chemical properties of two protein datasets and generates a model that can be used for further characterization (17). Briefly, the tool creates profiles, or physico-chemical signatures, for each protein in the K+ and Li+ binding sets, utilizing a set of 80 features - both experimentally and statistically derived from other algorithms available at https://tinyurl.com/yrbezujv. Specifically, we used hydrophobicity, alpha-helix, beta-sheet, disorder, burial, aggregation, membrane and nucleic acid-binding propensities. Only differentially enriched properties (p-values < 10−5 using Fisher’s exact test) were selected for the final model. The classification of chromatin proteins into foldend and unfolded G4A4 based on K+ and Li+ binding groups is available at https://tinyurl.com/2ywnk2a2. Further information can be found at http://s.tartaglialab.com/page/clever_suite.

### *cat*GRANULE predictions of phase separation

The tendency of proteins to phase separate is predicted through *cat*GRANULE (45). The algorithm exploits predictions of RNA binding ability and structural disordered propensities and was employed in our analysis to discriminate K+ and Li+ binding groups. Further information can be found at http://s.tartaglialab.com/new_submission/catGRANULE.

### G4 occurence predictions

G4 motifs predictions were carried out using *pqsfinder* package (version 2.8.0) in an R (4.1.0) enviroment. As input to the pipeline we used K+ and Li+ binding sites for Human protein–RNA interactions that were collected from eCLIP experiments (46) with stringent cut-offs [−log10(p value) > 3 and −log2(fold_enrichment) >3]. *pqsfinder* was used with default parameters and score = 52 (default) was used as score threshold score for accepting the occurence of a G4 motif.

### AlphaFold predictions of structural disorder

We used AlphaFold for calculations of protein structures (47). The PDBs, available from https://alphafold.ebi.ac.uk/, have been analyzed using STRIDE (48) to extract information on the structural content. The Coil, Turn, Helix and Strand content are reported in **Supplementary Table 5**.

## Notes

### Competing Interest Statement

The authors have declared no competing interest.

## References

1. Smith T, Villanueva E, Queiroz RML, Dawson CS, Elzek M, Urdaneta EC, et al. Organic phase separation opens up new opportunities to interrogate the RNA-binding proteome. Curr Opin Chem Biol. 2020 Feb;54:70–5.

2. Oksuz O, Henninger JE, Warneford-Thomson R, Zheng MM, Erb H, Overholt KJ, et al. Transcription factors interact with RNA to regulate genes. bioRxiv [Internet]. 2022; Available from: https://www.biorxiv.org/content/early/2022/09/28/2022.09.27.509776

3. Conrad T, Albrecht AS, Costa VRDM, Sauer S, Meierhofer D, Ørom UA. Serial interactome capture of the human cell nucleus. Nat Commun. 2016;

4. Baltz AG, Munschauer M, Schwanhäusser B, Vasile A, Murakawa Y, Schueler M, et al. The mRNA-Bound Proteome and Its Global Occupancy Profile on Protein-Coding Transcripts. Mol Cell. 2012;

5. Castello A, Fischer B, Eichelbaum K, Horos R, Beckmann BM, Strein C, et al. Insights into RNA Biology from an Atlas of Mammalian mRNA-Binding Proteins. Cell. 2012;

6. Perez-Perri JI, Ferring-Appel D, Huppertz I, Schwarzl T, Stein F, Rettel M, et al. The RNA-binding protein landscapes differ between mammalian organs and cultured cells. bioRxiv. 2022 Jan;2022.02.10.479897.

7. Bourdon S, Herviou P, Dumas L, Destefanis E, Zen A, Cammas A, et al. QUADRatlas: the RNA G-quadruplex and RG4-binding proteins database. Nucleic Acids Res. 2022 Sep 16;gkac782.

8. Quattrone A, Dassi E. The Architecture of the Human RNA-Binding Protein Regulatory Network. iScience. 2019 Nov 22;21:706–19.

9. Bhattacharyya D, Mirihana Arachchilage G, Basu S. Metal Cations in G-Quadruplex Folding and Stability. Front Chem. 2016;4:38.

10. Saldaña-Meyer R, Rodriguez-Hernaez J, Escobar T, Nishana M, Jácome-López K, Nora EP, et al. RNA Interactions Are Essential for CTCF-Mediated Genome Organization. Mol Cell. 2019 Nov;76(3):412-422.e5.

11. Li L, Williams P, Ren W, Wang MY, Gao Z, Miao W, et al. YY1 interacts with guanine quadruplexes to regulate DNA looping and gene expression. Nat Chem Biol. 2021 Feb;17(2):161–8.

12. Prado Martins R, Findakly S, Daskalogianni C, Teulade-Fichou MP, Blondel M, Fåhraeus R. In Cellulo Protein-mRNA Interaction Assay to Determine the Action of G-Quadruplex-Binding Molecules. Mol Basel Switz. 2018 Nov 29;23(12):E3124.

13. Bellucci M, Agostini F, Masin M, Tartaglia GG. Predicting protein associations with long noncoding RNAs. Nat Methods. 2011 Jun;8(6):444–5.

14. Cirillo D, Agostini F, Klus P, Marchese D, Rodriguez S, Bolognesi B, et al. Neurodegenerative diseases: quantitative predictions of protein-RNA interactions. RNA N Y N. 2013 Feb;19(2):129–40.

15. Fay MM, Lyons SM, Ivanov P. RNA G-Quadruplexes in Biology: Principles and Molecular Mechanisms. J Mol Biol. 2017 Jul 7;429(14):2127–47.

16. Agostini F, Zanzoni A, Klus P, Marchese D, Cirillo D, Tartaglia GG. CatRAPID omics: A web server for large-scale prediction of protein-RNA interactions. Bioinformatics. 2013;

17. Klus P, Bolognesi B, Agostini F, Marchese D, Zanzoni A, Tartaglia GG. The cleverSuite approach for protein characterization: predictions of structural properties, solubility, chaperone requirements and RNA-binding abilities. Bioinforma Oxf Engl. 2014 Jun 1;30(11):1601–8.

18. Castello A, Fischer B, Frese CK, Horos R, Alleaume AM, Foehr S, et al. Comprehensive Identification of RNA-Binding Domains in Human Cells. Mol Cell. 2016 Aug 18;63(4):696–710.

19. Radzicka A, Wolfenden R. Comparing the polarities of the amino acids: side-chain distribution coefficients between the vapor phase, cyclohexane, 1-octanol, and neutral aqueous solution. Biochemistry. 1988 Mar 8;27(5):1664–70.

20. Martin EW, Holehouse AS. Intrinsically disordered protein regions and phase separation: sequence determinants of assembly or lack thereof. Emerg Top Life Sci. 2020 Dec 11;4(3):307–29.

21. Zhang Y, Yang M, Duncan S, Yang X, Abdelhamid MAS, Huang L, et al. G-quadruplex structures trigger RNA phase separation. Nucleic Acids Res. 2019 Dec 16;47(22):11746–54.

22. Tsuruta M, Torii T, Kohata K, Kawauchi K, Tateishi-Karimata H, Sugimoto N, et al. Controlling liquid-liquid phase separation of G-quadruplex-forming RNAs in a sequencespecific manner. Chem Commun Camb Engl. 2022 Nov 22;58(93):12931–4.

23. Martadinata H, Phan AT. Structure of human telomeric RNA (TERRA): stacking of two G-quadruplex blocks in K(+) solution. Biochemistry. 2013 Apr 2;52(13):2176–83.

24. Asamitsu S, Yabuki Y, Matsuo K, Kawasaki M, Hirose Y, Kashiwazaki G, et al. RNA G-quadruplex organizes stress granule assembly through DNAPTP6 in neurons. Sci Adv. 2023 Feb 24;9(8):eade2035.

25. Han X, Yu D, Gu R, Jia Y, Wang Q, Jaganathan A, et al. Roles of the BRD4 short isoform in phase separation and active gene transcription. Nat Struct Mol Biol. 2020 Apr;27(4):333–41.

26. Chong S, Dugast-Darzacq C, Liu Z, Dong P, Dailey GM, Cattoglio C, et al. Imaging dynamic and selective low-complexity domain interactions that control gene transcription. Science. 2018 Jul 27;361(6400):eaar2555.

27. Kumari S, Bugaut A, Huppert JL, Balasubramanian S. An RNA G-quadruplex in the 5’ UTR of the NRAS proto-oncogene modulates translation. Nat Chem Biol. 2007 Apr;3(4):218–21.

28. Morris MJ, Negishi Y, Pazsint C, Schonhoft JD, Basu S. An RNA G-quadruplex is essential for cap-independent translation initiation in human VEGF IRES. J Am Chem Soc. 2010 Dec 22;132(50):17831–9.

29. Kwok CK, Marsico G, Sahakyan AB, Chambers VS, Balasubramanian S. rG4-seq reveals widespread formation of G-quadruplex structures in the human transcriptome. Nat Methods. 2016 Oct;13(10):841–4.

30. Shahid R, Bugaut A, Balasubramanian S. The BCL-2 5’ untranslated region contains an RNA G-quadruplex-forming motif that modulates protein expression. Biochemistry. 2010 Sep 28;49(38):8300–6.

31. Kharel P, Becker G, Tsvetkov V, Ivanov P. Properties and biological impact of RNA G-quadruplexes: from order to turmoil and back. Nucleic Acids Res. 2020 Dec 16;48(22):12534–55.

32. Kharel P, Fay M, Manasova EV, Anderson PJ, Kurkin AV, Guo JU, et al. Stress promotes RNA G-quadruplex folding in human cells. Nat Commun. 2023 Jan 13;14(1):205.

33. Sauer M, Juranek SA, Marks J, De Magis A, Kazemier HG, Hilbig D, et al. DHX36 prevents the accumulation of translationally inactive mRNAs with G4-structures in untranslated regions. Nat Commun. 2019 Jun 3;10(1):2421.

34. Chong PA, Vernon RM, Forman-Kay JD. RGG/RG Motif Regions in RNA Binding and Phase Separation. J Mol Biol. 2018 Nov 2;430(23):4650–65.

35. Youn JY, Dunham WH, Hong SJ, Knight JDR, Bashkurov M, Chen GI, et al. High-Density Proximity Mapping Reveals the Subcellular Organization of mRNA-Associated Granules and Bodies. Mol Cell. 2018 Feb 1;69(3):517-532.e11.

36. Hondele M, Sachdev R, Heinrich S, Wang J, Vallotton P, Fontoura BMA, et al. DEAD-box ATPases are global regulators of phase-separated organelles. Nature. 2019 Sep;573(7772):144–8.

37. Hou Y, Li F, Zhang R, Li S, Liu H, Qin ZS, et al. Integrative characterization of G-Quadruplexes in the three-dimensional chromatin structure. Epigenetics. 2019 Sep;14(9):894–911.

38. Keller JG, Hymøller KM, Thorsager ME, Hansen NY, Erlandsen JU, Tesauro C, et al. Topoisomerase 1 inhibits MYC promoter activity by inducing G-quadruplex formation. Nucleic Acids Res. 2022 Jun 24;50(11):6332–42.

39. Sørensen CS, Hansen LT, Dziegielewski J, Syljuåsen RG, Lundin C, Bartek J, et al. The cell-cycle checkpoint kinase Chk1 is required for mammalian homologous recombination repair. Nat Cell Biol. 2005 Feb;7(2):195–201.

40. Bader AS, Hawley BR, Wilczynska A, Bushell M. The roles of RNA in DNA double-strand break repair. Br J Cancer. 2020 Mar;122(5):613–23.

41. Santos A, Colaço AR, Nielsen AB, Niu L, Geyer PE, Coscia F, et al. Clinical Knowledge Graph Integrates Proteomics Data into Clinical Decision-Making [Internet]. Bioinformatics; 2020 May [cited 2023 Mar 27]. Available from: http://biorxiv.org/lookup/doi/10.1101/2020.05.09.084897

42. Lazar C, Gatto L, Ferro M, Bruley C, Burger T. Accounting for the Multiple Natures of Missing Values in Label-Free Quantitative Proteomics Data Sets to Compare Imputation Strategies. J Proteome Res. 2016 Apr 1;15(4):1116–25.

43. Cirillo D, Blanco M, Armaos A, Buness A, Avner P, Guttman M, et al. Quantitative predictions of protein interactions with long noncoding RNAs. Nat Methods. 2016 Dec 29;14(1):5–6.

44. Lang B, Armaos A, Tartaglia GG. RNAct: Protein-RNA interaction predictions for model organisms with supporting experimental data. Nucleic Acids Res. 2019 Jan 8;47(D1):D601–6.

45. Bolognesi B, Lorenzo Gotor N, Dhar R, Cirillo D, Baldrighi M, Tartaglia GG, et al. A Concentration-Dependent Liquid Phase Separation Can Cause Toxicity upon Increased Protein Expression. Cell Rep. 2016 Jun 28;16(1):222–31.

46. Van Nostrand EL, Freese P, Pratt GA, Wang X, Wei X, Xiao R, et al. A large-scale binding and functional map of human RNA-binding proteins. Nature. 2020 Jul;583(7818):711–9.

47. Jumper J, Evans R, Pritzel A, Green T, Figurnov M, Ronneberger O, et al. Highly accurate protein structure prediction with AlphaFold. Nature. 2021 Aug;596(7873):583–9.

48. Heinig M, Frishman D. STRIDE: a web server for secondary structure assignment from known atomic coordinates of proteins. Nucleic Acids Res. 2004 Jul 1;32(Web Server issue):W500-502.

