## Supplementary figures and images for "Identification and prediction of G-quadruplex RNA-binding proteins with roles in transcription and phase separation"

### Supplementary Figure 1

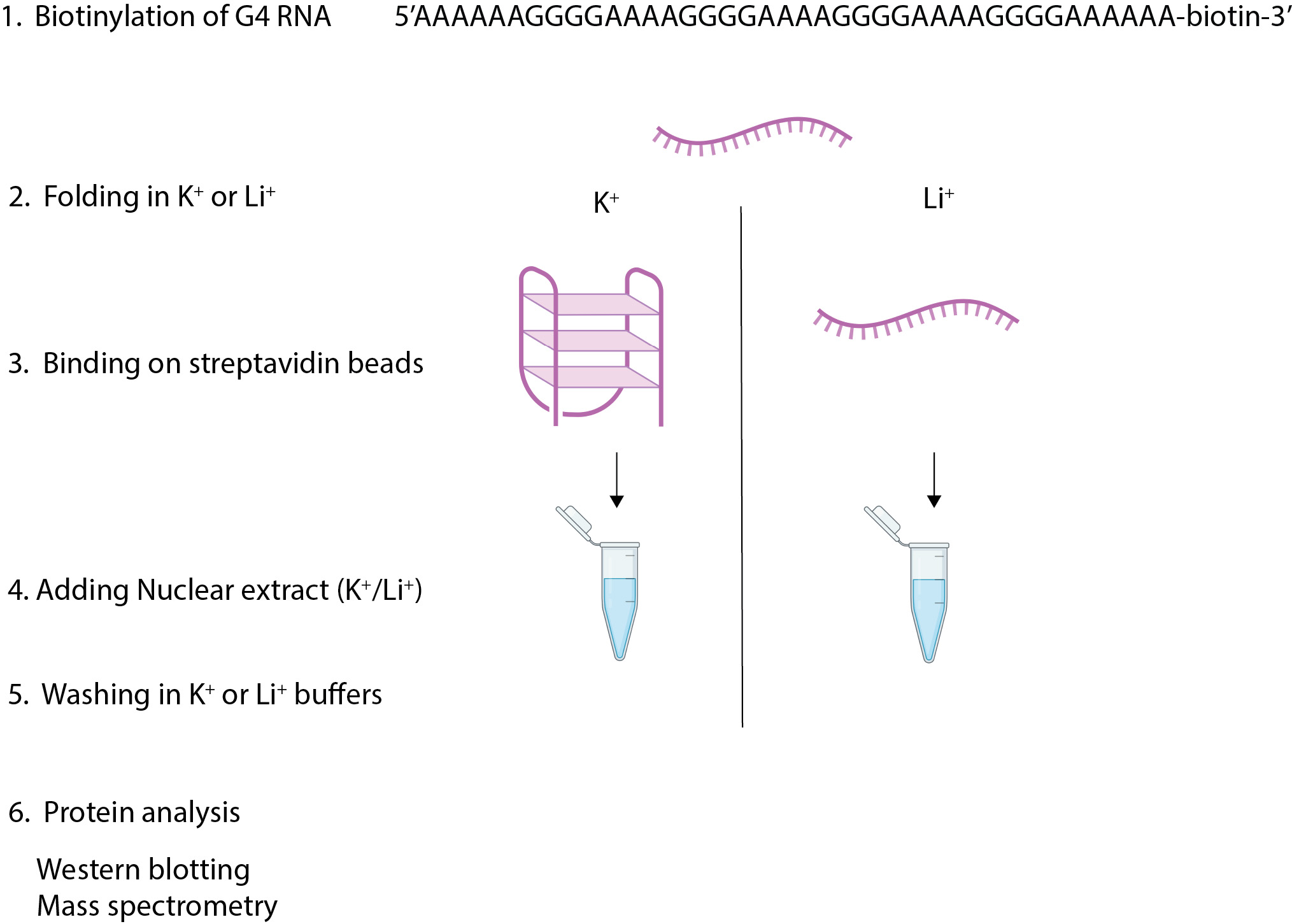
